# Generating quantitative binding landscapes through fractional binding selections, deep sequencing and data normalization

**DOI:** 10.1101/509984

**Authors:** Michael Heyne, Niv Papo, Julia Shifman

## Abstract

Quantifying the effects of various mutations on binding free energy is crucial for understanding the evolution of protein-protein interactions and would greatly facilitate protein engineering studies. Yet, measuring changes in binding free energy (ΔΔG_bind_) remains a tedious task that requires expression of each mutant, its purification, and affinity measurements. We developed a new approach that allows us to quantify ΔΔG_bind_ for thousands of protein mutants in one experiment. Our protocol combines protein randomization, Yeast Surface Display technology, Next Generation Sequencing, and a few experimental ΔΔG_bind_ data points on purified proteins to generate ΔΔG_bind_ values for the remaining numerous mutants of the same protein complex. Using this methodology, we comprehensively map the single-mutant binding landscape of one of the highest-affinity interaction between BPTI and Bovine Trypsin. We show that ΔΔG_bind_ for this interaction could be quantified with high accuracy over the range of 12 kcal/mol displayed by various BPTI single mutants.

## Introduction

Protein-protein interactions (PPIs) control virtually all processes in the cell. Mutations at PPI binding interfaces frequently affect free energy of binding (ΔΔG_bind_), sometimes abrogating and sometimes stabilizing the interaction. This change in binding affinity of one PPI could translate into remodeling of the whole PPI network, frequently leading to dysregulation of signal transduction pathways and disease^1,2^. Therefore, understanding how specific mutations in protein complexes affect their binding affinity is extremely important to both basic biology and to biomedical sciences, where inhibition of a particular PPI might help to treat the related disease.

In the recent years, many groups reported computational methods for predicting ΔΔG_bind_ from structure and/or sequence^3-9^. While achieving good predictions on average, these methods frequently give large errors in particular cases, revealing that our comprehension of the precise molecular forces that govern binding affinity in PPIs remains incomplete^10^. Our knowledge in this area could be greatly expanded by acquiring large sets of data for ΔΔG_bind_ values in various PPIs, facilitating progress in computational modeling. Yet, experiments that determine ΔΔG_bind_ remain laborious since they involve DNA manipulation, protein expression and purification in different organisms and binding affinity measurements using different techniques. Thus, experimental data describing mutational effects on binding affinity for each particular PPI remain sparse and sometimes inconsistent between different reported experiments^11^. Furthermore, the majority of mutations reported in the literature are mutations to alanine^11-14^. Such mutations are of limited use for studies of protein evolution and protein engineering since they most frequently lead to complex destabilization and are rarely observed in nature.

A much more attractive and informative approach would be to explore all possible mutational effects for a particular PPI in a single experiment, thus generating a comprehensive binding landscape for this PPI^15,16^. Such binding landscapes could be used to define evolutionary paths accessible to a particular PPI, to characterize energetic contribution of each position, and to locate frequently sought affinity- and specificity-enhancing mutations^4,17^. First efforts in this direction utilized phage display technology that allows to select binders from a large combinatorial library of protein mutants^16,18,19^. Through several rounds of selection, protein mutants compatible with binding to a particular target are selected. Subsequent sequencing of multiple selected clones allows us to calculate the frequency of each amino acid at each position, providing information on binding hot-spots^20^ and cold-spots^21^. Further studies in this direction utilized Yeast Surface Display (YSD)^22^ for selecting protein binders coupled to Next-Generation sequencing (NGS) to produce binding landscapes for various PPIs^23^. While YSD enables fast sorting using Fluorescently Activated Cell Sorting (FACS), NGS permits more accurate calculation of amino acid frequencies for each of the detected mutants. The ratio between the amino acid frequency in the selected pool of binders and the same frequency in the initial naïve library, referred to as the enrichment value, is calculated for each amino acid at each of the explored position. The enrichment values are then plotted to produce PPI binding landscapes. In such an approach, Whitehead *et al.* mapped the full single-mutant binding landscape of a *de novo* designed inhibitor interacting with hemagglutinin and used the enrichment information to design affinity-matured inhibitors^24^.

In spite of great promise of this approach, further studies on different biological systems revealed its potential limitations. While affinity enhancing mutations could be readily identified by this methodology, relatively low correlation (R value of 0.5) between the NGS-derived enrichment values and experimental ΔΔG_bind_ values for purified proteins was observed^17^. Additional studies showed that ΔΔG_bind_ could be inferred from the NGS-based enrichment values only in the narrow range of energies from −0.8 to +0.5 kcal/mol^25,26^, preventing construction of quantitative binding landscapes for all of the explored mutations with broader range of target affinities. In addition, the above methodology set a requirement on the concentration of the target protein in the selection experiment; the concentration should be similar to the interaction K_d_, thus limiting the application of the approach to only subset of all PPIs with medium affinities. For high-affinity PPIs (Kd < 10^−10^ M), this condition would imply the usage of very low target protein concentration, resulting in undetectable fluorescence signal by FACS. For low-affinity PPIs (Kd > 10^−5^ M), high concentrations of protein would be necessary, making such experiments impractical.

We introduce a novel approach that allows us to overcome the abovementioned limitations and to generate quantitative binding landscapes for any PPI, regardless of their K_d_ value. Here, we demonstrate the applicability of our approach to a particularly difficult target, a complex between Bovine Trypsin (BT) and its inhibitor BPTI that possesses ultra-high affinity of 10^−14^ M. We show that through our high-throughput NGS-based approach, we can obtain ΔΔG_bind_ values for all BPTI binding interface mutants that correlate extremely well with experimental results on purified proteins over the range of more than 12 kcal/mol free energy changes. Our method allowed us to comprehensively map the binding landscape for this ultra-high affinity interaction, which would be impossible using any alternative technique.

## Results and Discussion

To demonstrate how our approach could be used to produce quantitative binding landscapes, we first prepared the BPTI/BT complex for YSD experiments. For this purpose, the BPTI WT gene was incorporated into the pCTCON vector, that facilitates BPTI expression on the surface of yeast cells with a C-terminal myc-tag (cMyc) for monitoring protein expression (**Figure 1A**). Binding of BT to BPTI mutants was accessed by monitoring fluorescence of the FITC fluorophore conjugated to a biotinylated BT via NeutrAvidin. The assessment of binding of BPTIwt to BT by FACS showed a diagonal narrow distribution, demonstrating that BPTI is well expressed on the surface of yeast cells, is properly folded, and binds to BT (**Figure S1**).

**Figure 1:**
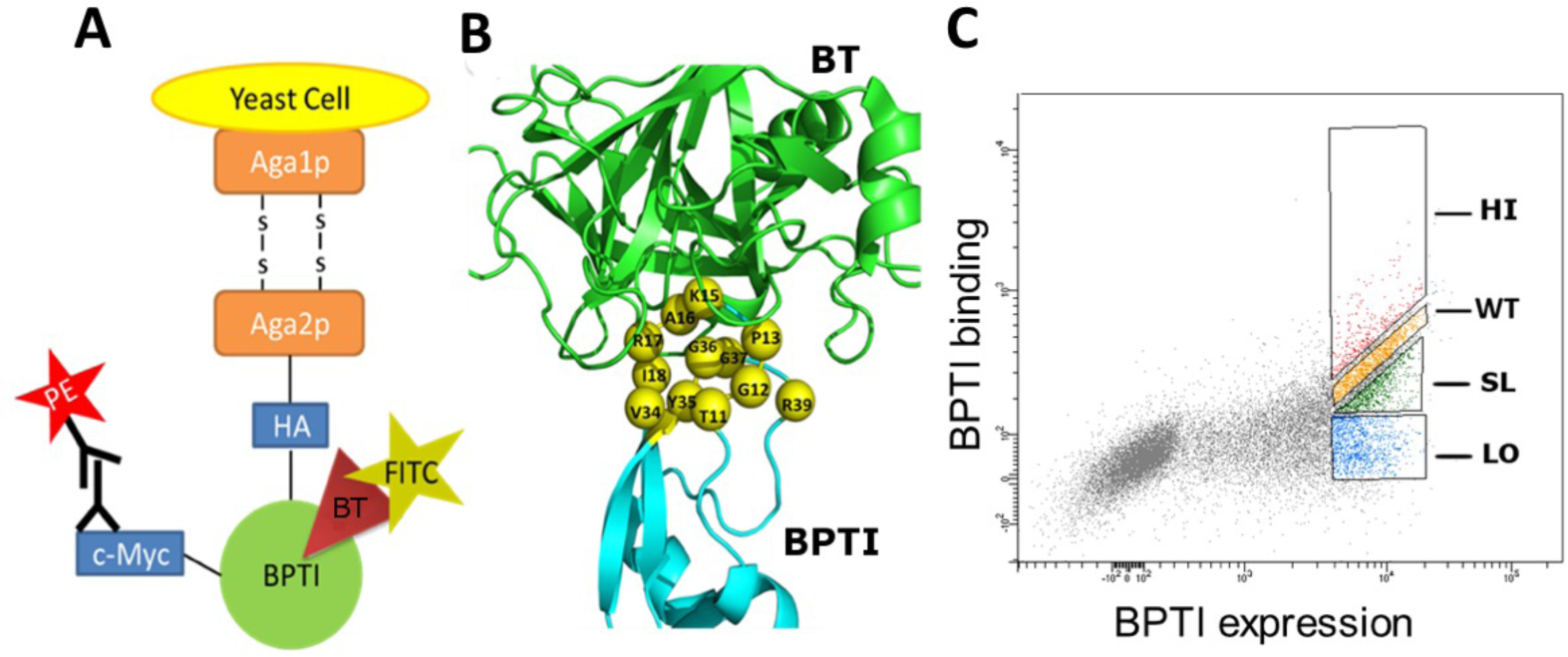
A) Yeast surface display construct with BPTI displayed on the surface of yeast cells B) Construction of the BPTI single mutant library. Structure of the BT/BPTI complex as shown from PDB 3OTJ. BT is colored in green, BPTI is colored in cyan and the BPTI binding interface positions are shown as spheres, labelled and colored in yellow. (C) FACS data showing sorting of four different populations of the BT/BPTI complex. The PE signal monitoring BPTI expression is shown on the x-axis and the FITC signal monitoring binding between fluorescently-labelled BT and BPTI variants expressed on the yeast surface is shown on the y-axis. The uppermost sorting gate HI represents all mutants with affinity higher than WT. The second uppermost gate WP represents all mutants with an affinity similar to BPTI_WT_. The third gate SL represents all mutants with an affinity slightly lower than WT and the undermost gate LO represents all mutants with an affinity much lower than WT. BT concentration was 5 nM in this experiment.

Next, a combinatorial library was generated containing all single BPTI mutants at positions that are in the direct binding interface with BT. Thus, twelve BPTI positions were randomized to all twenty amino acids with an NNS codon, leaving intact two cysteines (C14 and C38) that form a disulfide bond and thus are crucial for BPTI folding (**Figure 1B**). The library of 228 (19 x 12) BPTI single mutants was constructed using the TPCR protocol^27^. The BPTI mutant library was expressed on the surface of yeast cells and incubated with a fluorescently labeled BT at concentration of 5 nM. This concentration of BT, although five orders of magnitude higher than the K_d_ of BT/BPTI interaction, was chosen since it was the minimum concentration of BT that resulted in a considerable spread of FACS binding signals from different BPTI mutants to BT (**Figure 1C**).

### Improving accuracy and extending prediction range by collecting more data

One of the limitations of previous approaches for binding landscape generation was that ΔΔG_bind_ showed linear dependence on the NGS-enrichment value only in the narrow range of ΔΔG_bind_ values close to zero^26^. The methodology could not previously discriminate between different highly destabilizing mutations since such mutations were characterized with the same enrichment values. The same was true for mutations that showed high stabilization of the complex. To overcome this limitation and to increase the range of sensitivity for ΔΔG_bind_ predictions, we introduced our first innovation and used multiple affinity gates from which the mutants were collected during the YSD selection experiment. The multiple gates would allow us to collect information for each mutant several times, and each particular mutant would be enriched in at least one affinity gate. In this particular work, we used four affinity gates for mutant collection: higher than WT affinity (HI), WT-like affinity (WT), slightly lower than WT affinity (SL), and strongly lower than WT affinity (LO) (**Figure 1C**). The WT affinity gate was set according to the FACS signal produced by BPTIwt binding to BT at same conditions (**Figure 1S**). The cells from each gate were then grown, analyzed for binding to BT (**Figure S4**) and sequenced with NGS, resulting in 300- 900K reads per each population. In addition, the naïve pre-sorted library of BPTI mutants was sequenced.

We further assessed the quality of the NGS data using synonymous mutations as a test. Since some errors in the data could come from errors in the NGS process, especially for sequences detected with low frequency, we tested different cut-off values below which the data on the BPTI mutant would be discarded. Using different cut-off values, we calculated deviations in enrichment values for synonymous mutations expressing the same BPTI variant. Our data shows that at the cutoff value of 100 sequences per BPTI mutant, deviations in enrichment values were negligible (< 0.001) (**Figure S2**). Using this threshold, we were able to detect all 228 BPTI single mutants present in the naïve library. No threshold was applied to the sorted populations, since in such population the low number of sequences was caused by the depletion of that mutant from the population.

We thus had in our hands four enrichment values from four affinity gates for each of the 228 BPTI mutants (**Figure 2**). Closer examination of the data showed that enrichment values in HI and LO affinity gates exhibited pseudo-symmetry, with highly enriched mutations in the HI gate being highly depleted in the LO gate and vice versa. The enrichment value maps could be used to define binding hot-spots for the BPTI/BT interactions (such as position 15, 16 indicated as red starts on top of **Figure 2**) and more tolerant to mutations positions (such as 11 and 34 indicated as blue stars on **Figure 2**). However, these maps were not sufficient in determining exact ΔΔG_bind_ values for each of the mutation. In fact, we noticed that some mutations that showed enrichment value of ~1 in the HI gate, that should correspond to the WT-like affinity, were determined to be destabilizing when measured with purified proteins (for example, G12A with experimental ΔΔG_bind_ of +4.35 kcal/mol^28-31^). This over-prediction of neutral and affinity-enhancing mutations by our NGS results was due to the fact that in the YSD selection experiment we used much higher concentration of BT compared to the K_d_ of BPTI/BT interaction, shifting the equilibrium towards protein binding even for those BPTI mutants that possessed weaker affinities compared to the WT protein. To overcome this problem, we introduced the second innovation, described below.

**Figure 2:**
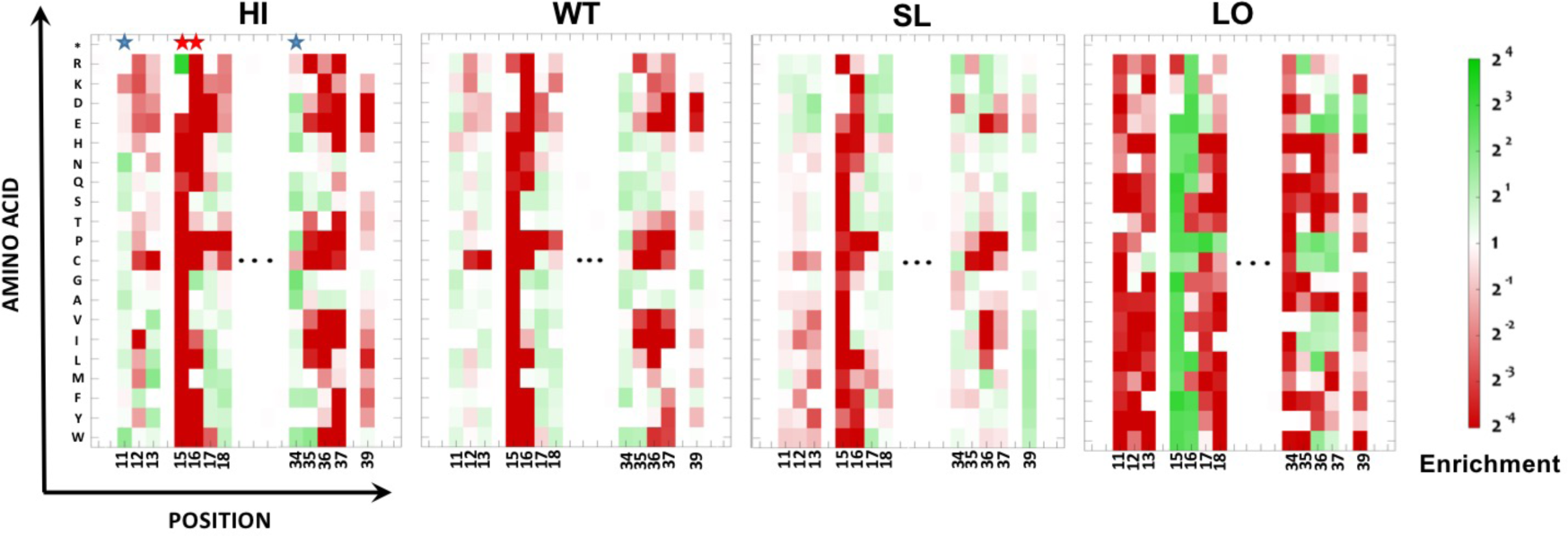
Experimental Binding landscapes for the BT/BPTI complex. These heat maps show the enrichment of each variant in each of the four sorted affinity gates relative to the pre-sort. The enrichment ratio varies from high (green) to low (red) as shown on the right axis. Red and blue stars on top show binding hot-spots and cold spots, respectively. Similar maps were obtained for all four PPIs.

### Normalizing NGS data to get quantitative ΔΔG_bind_ measurements

Our second innovation was to obtain a normalization formula for converting the enrichment data from four affinity gates into one ΔΔG_bind_ value. For this purpose, we collected all available ΔΔG_bind_ experimental data for binding of BPTI single mutants to BT, comprising 29 data points^28-31^. Plotting the experimental ΔΔG_bind_ vs. enrichment values for each of the four affinity gates showed that ΔΔG_bind_ was linearly dependent on the natural log of enrichment values in HI and LO gates (R-value of 0.87 for each of the gates, **Figure S3**). The NGS values from HI and LO gates were denoted further as functions X1 and X4, respectively. ΔΔG_bind_ showed a more complicated two-valued function behavior for WT and SL gates. This was expected since for these gates, the highest enrichment values were observed in the narrow range of ΔΔG_bind_ values but decreased for mutations that showed both higher and lower affinities compared to that narrow range of values. To eliminate this complicated multi-variable behavior and at the same time to utilize the additional information from WT and SL gates, we constructed two additional functions that multiplied enrichment values from HI and SL gates (HI x SL, denoted further as X2) and enrichment values from WT and LO gates (WT x LO, denoted further as X3). These two functions, X2 and X3, were linearly dependent on ΔΔG_bind_. We next used a linear regression to produce the best possible fit of experimental ΔΔG_bind_ values using the linear combination of four functions (X1, X2, X3, X4) arriving to the normalization formula that converts NGS enrichment values into ΔΔG_bind_ for the BPTI/BT complex in this particular experiment:

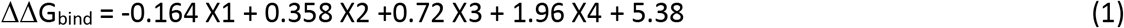

Using 29 data points and leave-one-out cross-validation approach, where each data point was predicted without the enrichment information for that particular data point, we were able to predict experimental ΔΔG_bind_ values with very high accuracy over the range of more than 12 kcal/mol (**Figure 3**; R = 0.90, σ =1.5 kcal/mol). We used the obtained normalization formula to convert the enrichment values to ΔΔG_bind_ values for all the single BPTI mutants in the library. Since the enrichment values for these mutants are within the range explored in our small experimental data set, the calculated ΔΔG_bind_ values are expected to have the same accuracy as reported above.

**Figure 3:**
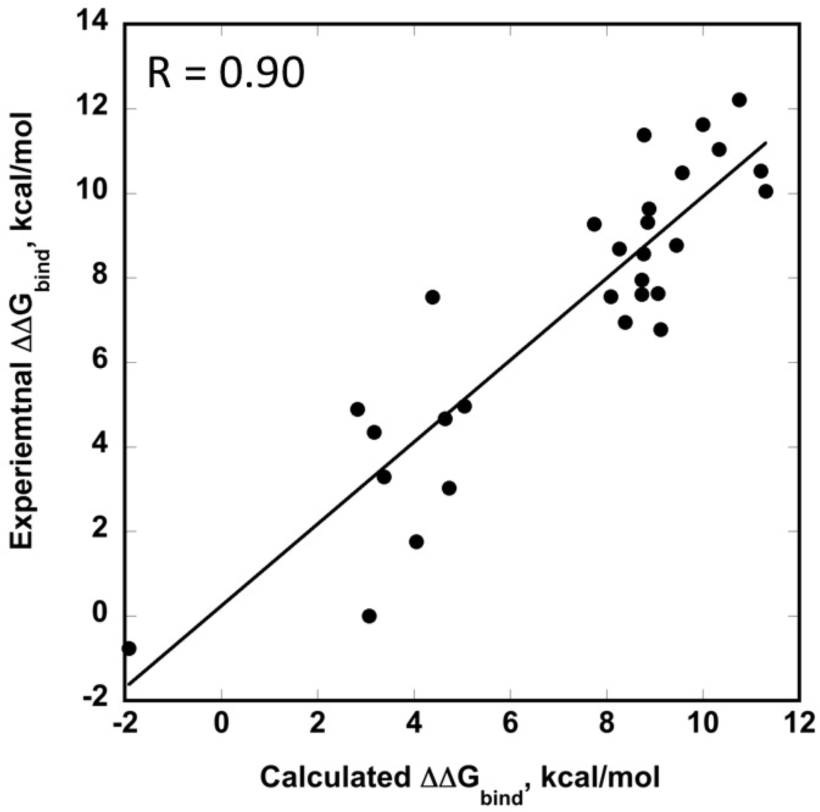
Correlation between the calculated ΔΔG_bind_ using the enrichment values from the NGS results and the ΔΔG_bind_ measurements for purified protein variants of BPTI interacting with BT. Leave-one-out cross validation was used to produce this graph.

**Figure 4** shows the ΔΔG_bind_ values for all single BPTI mutants at 12 binding interface positions interacting with BT, producing a quantitative binding landscape for this PPI. As can be seen, the majority of the mutations at all positions in this PPI are highly destabilizing, producing destabilization as high as almost 12 kcal/mol. The most non-tolerant to substitution positions are 15 and 16 that lie in the core of the binding interface (**Figure 1B**). However, at position 15, one mutation, K15R, was determined to substantially stabilize the complex, in agreement with experimental results on purified proteins^30^. **Figure 4** also shows that the same type of mutations (e. g. hydrophobic or polar) frequently produce similar changes in ΔΔG_bind_ for the same position. We thus established that the BPTI/BT complex with the K_d_ of 10^−14^ M is highly optimized by nature, with most single mutations in BPTI leading to high destabilization of the PPI, and very few neutral and affinity-enhancing mutations.

**Figure 4:**
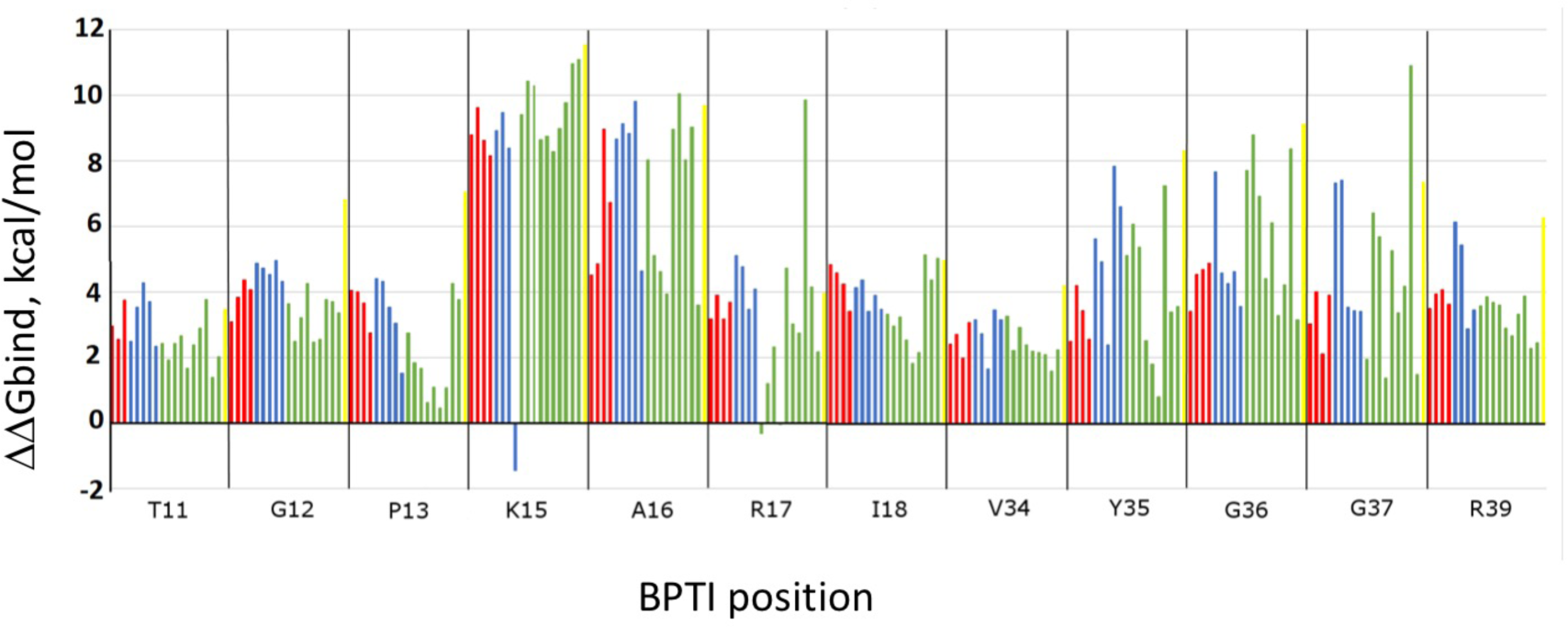
Changes in ΔΔG_bind_ for all single mutants of BPTI interacting with BT. Each bar represents a mutation to one amino acid including hydrophobic amino acids (green), polar amino acids (red), charged amino acids (blue) and Cys (yellow). X-axis shows WT residue followed by position.

In summary, we report a novel approach that allows us for the first time to produce quantitative binding landscapes based on binding selections into several affinity gates, NGS of the selected mutants, and normalization procedure using a small data set of experimentally determined ΔΔG_bind_ values. Very recently, a similar direction has been taken by Keating and colleagues to design mutations and improve affinity in peptide/protein interactions^32^. Unlike our study, the authors used IC50 value of yeast titrations to normalize NGS data and to predict ΔΔG_bind_ for various peptide mutants with K_d_s in the medium affinity range. A similar thinking by two independent studies attest to attractiveness of our methodology for binding landscape mapping. Yet our work presents a simpler normalization procedure based on actual *in vitro* affinity measurements and allows to explore PPIs at the limits of the Kd spectra. We demonstrate superior correlations between ΔΔG_bind_ predictions and actual *in vitro* measurements over a much larger range compared to all previously reported approaches.

To achieve good prediction accuracy, the experimental data points used for normalization should show large spread in ΔΔG_bind_ values, including both affinity-enhancing and affinity-reducing mutations. More affinity gates could be used in future experiments, although in this case normalization would require a larger set of experimental data; at least five data points per each parameter should be used in the normalization function to avoid overfitting. Our methodology could be applied to study the evolutionary paths of any PPI regardless of its K_d_ value and to compare binding landscapes of various PPIs. The approach could be easily extended to studies of double and higher-order mutational steps, providing more comprehensive information on PPI evolution and facilitating new modeling and protein engineering studies.

## Acknowledgements

We thank I. Cohen and B. Gazit for help with initial experiments and data analysis and Y. Peleg for help with molecular biology experiments. We also thank A. Zilka for help with FACS experiments. This work was supported by the Israel Science Foundation (ISF) grant 1873/15 (J. M. S.) and by the European Research Council (ERC) grant 336041 and the ISF grant 615/14 (N. P.)

