## Supplementary Information for "Generating quantitative binding landscapes through fractional binding selections, deep sequencing and data normalization"

### Methods

#### BPTI library construction

The $BPTI_{WT}$ was generated by PCR using overlapping oligonucleotides. The final PCR assembled fragment was gel-purified and cloned into pCTCON vector via transformation by electroporation of EBY100 yeast cells and homologous recombination with the linearized vector (digested with *Nhe*I and *Bam*HI), as previously described[1]. Twelve BPTI libraries were constructed from the $BPTI_{WT}$ gene by randomizing each of the binding interface positions with an NNS codon utilizing a TPCR protocol with one forward and one backward primer[2,3] . The PCR product was treated with DpnI to remove any parental plasmid, cleaned up with magnetic beads, transformed into *E. coli* and selected colonies were sequenced to confirm the successful generation and transformation of the BPTI library. The DNA containing each BPTI library was extracted and all the sublibraries were pooled together and balanced by their DNA concertation. Then, the pooled naïve library of BPTI single mutants was transferred into *S. cerevisiae* using 20 transformations resulting into 60,000 – 70,000 colonies for the complete library.

#### YSD sorting experiments

Yeast cells displaying the BPTI library or the $BPTI_{WT}$ on the YSD were grown in SDCAA selective medium and induced for BPTI protein expression with a galactose-containing SGCAA medium (as for SDCAA, but with galactose instead of dextrose), as previously described[4]. BPTI expression and binding to BT was measured by phycoerythrin and NeutrAvidin in dual-color flow cytometry (Accuri C6, BD Biosciences), respectively. The yeast cells were next sorted into four populations by FACSAria (BD Biosciences, San Jose, CA) including HI, WP, SL, and LO populations. Sorted cells were then grown in a selective medium, the plasmidic DNA was extracted for each of the sorted population and the naïve library and submitted to NGS by MiSeq, Illumina (service provided by Hylabs, Rechovot, IL).

#### NGS analysis

The paired-end reads from the NGS experiments were merged[5] and their quality scores were calculated in the FastQC tool[6]. In the Matlab script, the sequences were aligned, and sequences containing more than one mutation were filtered out. The number of each remaining BPTI mutation i in position j was counted in the sorted and the naïve populations and its frequency $f^{i,j}$ in the libraries was calculated (**Eq. 1**). Using the frequency of the mutant in one of the sorted populations and the naïve population, the enrichment $e^{i,j}$ of each BPTI mutant was calculated (**Eq**. **2**).

$$f^{i,j} = \frac{count^{i,j}}{count_{total}} \qquad (1)$$

$$e^{i,j} = \ln\left(\frac{(f^{i,j})_{sorted}}{(f^{i,j})_{naïve}}\right) \qquad (2)$$

All available experimental data on $\Delta\Delta G_{\text{bind}}$ for the BPTI/BT complex was used to obtain the best normalization formula for converting enrichment values from four sorted populations into $\Delta\Delta G_{\text{bind}}$ values. For this we used a linear regression model function in Mathematica (Wolfram Research) with 5 parameters as described in the manuscript. This normalization formula was further used to calculate $\Delta\Delta G_{\text{bind}}$ values for the remaining single BPTI mutants, for which no $\Delta\Delta G_{\text{bind}}$ values were previously measured.

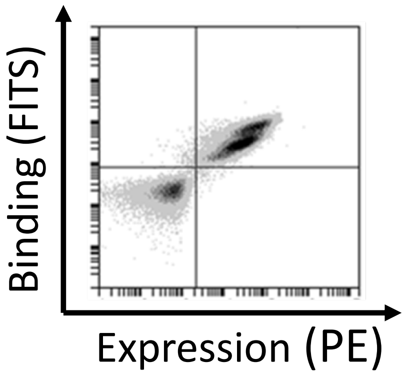


**Supplementary Figure S1:** FACS data of $BPTI_{WT}$ binding to 5 nM BT. BPTI Expression was monitored by PE fluorescence while binding to BT was monitored by FITS conjugated to BT.

A

B

C

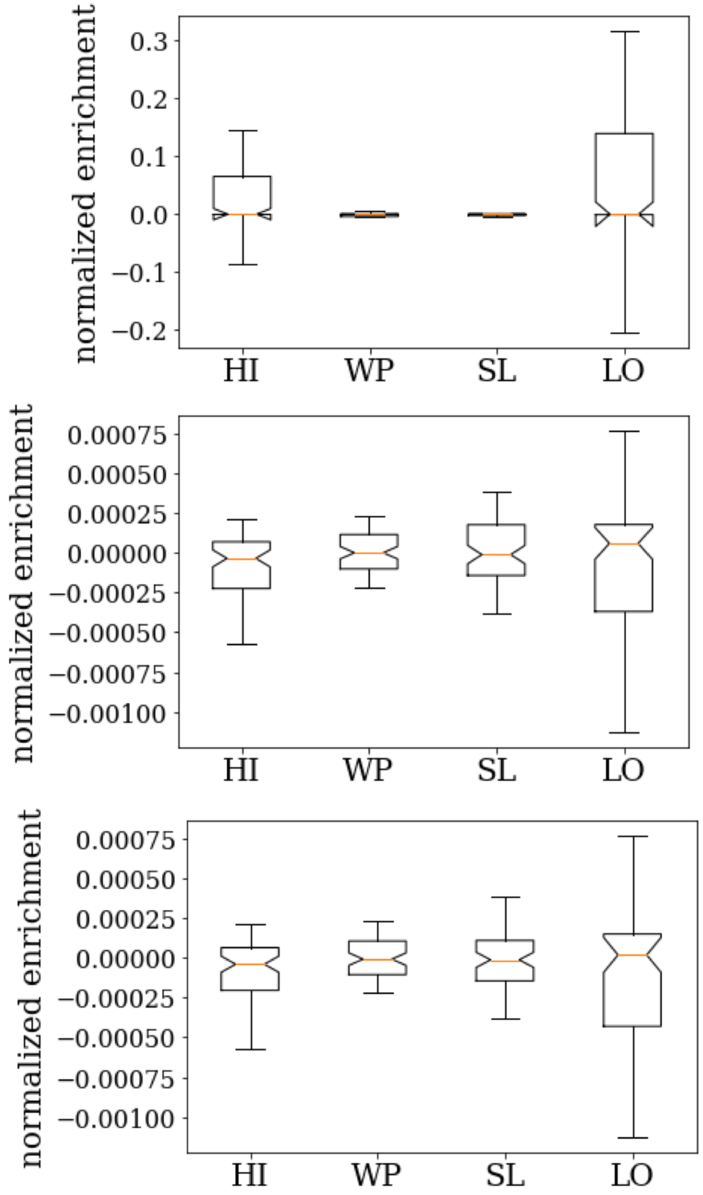


**Supplementary Figure S2:** Boxplots showing the distribution of the log2 of enrichment of synonymous mutations normalized to the enrichment of the DNA WT sequence (outliers are removed for better visualization) for three different cutoff thresholds: A) 10, B) 100 and C) 250.

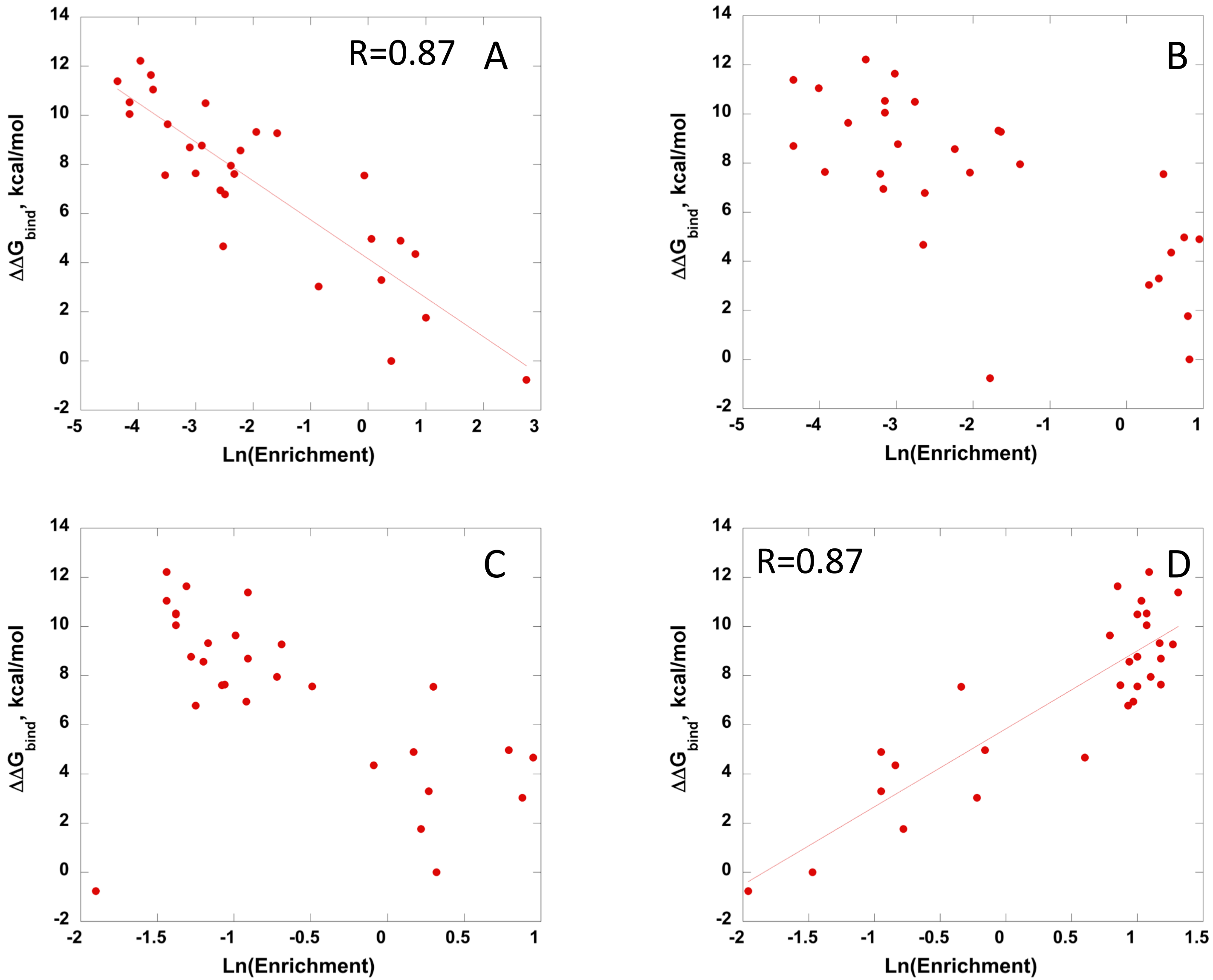


**Supplementary Figure S3:** Dependence of experimental $\Delta\Delta G_{bind}$ values on enrichment value from each gate: (A) HI; (B) WT; (C) SL; (D) LO.

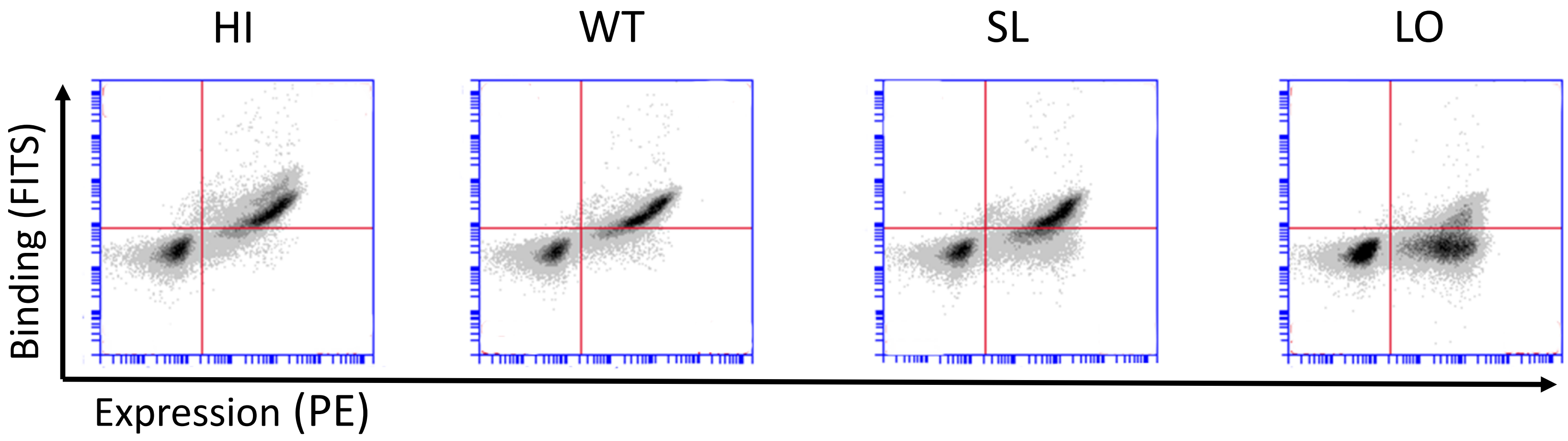


**Supplementary Figure S4:** Separation of BPTI mutant clones into four affinity windows: HI, WT, SL, and LO. FACS data after sorting with 5 nM BT is shown.
